# The SLIM1 transcription factor is required for arsenic resistance in *Arabidopsis thaliana*

**DOI:** 10.1101/2021.01.12.426316

**Authors:** Timothy O. Jobe, Qi Yu, Felix Hauser, Qingqing Xie, Yuan Meng, Tim Maassen, Stanislav Kopriva, Julian I. Schroeder

## Abstract

The transcriptional regulators of arsenic-induced gene expression remain largely unknown. Sulfur assimilation is tightly linked with arsenic detoxification. Here we report that mutant alleles in the SLIM1 transcription factor are substantially more sensitive to arsenic than cadmium. Arsenic treatment caused high levels of oxidative stress in the *slim1* mutants, and *slim1* alleles were impaired in both thiol and sulfate accumulation. We further found enhanced arsenic accumulation in roots of *slim1* mutants. Transcriptome analyses indicate an important role for SLIM1 in arsenic-induced tolerance mechanisms. The present study identifies the SLIM1 transcription factor as an essential component in arsenic tolerance and arsenic-induced gene expression. Our results suggest that the severe arsenic sensitivity of the *slim1* mutants is caused by altered redox status.

## Introduction

Many advanced technologies used by modern society rely on heavy metals and arsenic. These elements are toxic and pose a significant risk to the environment and human health if consumed. However, unlike animals, plants are often partially tolerant to heavy metals and arsenic and can accumulate large amounts in diverse tissues [1]. Arsenic is a highly toxic substance commonly found in anthropogenic wastes (electronics and fertilizers) and can also be found at high levels in certain rocks, soils, and waters globally [2–5]. While this toxic metalloid has no recognized role in plant or animal nutrition, plant-based products are the main entry point for arsenic into the food chain [6]. Thus, understanding the molecular mechanisms underlying plant uptake, transport, detoxification, and accumulation of arsenic is vital for enhancing the nutritional value and safety of our food.

We previously described the development of a plant genetic reporter line that fused the promoter of a cadmium and arsenic-inducible high-affinity sulfate transporter to firefly luciferase (*pSULTR1;2::LUC*) to identify mutants in signaling [7]. A major goal of this work was to identify the transcriptional regulators mediating rapid arsenic-induced gene expression in Arabidopsis. This approach was successful in identifying new alleles of the glutathione biosynthesis genes gamma-glutamylcysteine synthetase (γ-ECS) and glutathione synthetase (GS), as being required for cadmium and arsenic-induced gene expression [7]. Glutathione is necessary for the synthesis of phytochelatins, which detoxify many toxic compounds, including cadmium and arsenic, by chelation and sequestration in the vacuole[1,8–11]. Phytochelatins are short polymers of glutathione synthesized in the cytosol in response to toxic metal(loid)s. Thus, arsenic exposure can rapidly deplete glutathione levels, creating a high demand for glutathione in plant cells.

Because the tripeptide glutathione (Glu-Cys-Gly) contains the sulfur-containing amino acid cysteine, the sulfate assimilation pathway is inextricably linked to glutathione biosynthesis. Sulfate assimilation takes oxidized sulfur in the form of sulfate and, through a series of energy-dependent reducing steps, produces sulfide. Due to the toxicity of sulfide, this intermediate quickly reacts with O-acetylserine to produce the amino acid cysteine[12]. Thus, unlike animals, plants do not require exogenous sulfur-containing amino acids and proteins for survival[13]. More importantly, this creates a direct link between the sulfate assimilation pathway and the ability of plants to detoxify arsenic.

While our luciferase genetic reporter approach has not identified transcriptional regulators of arsenic-induced gene induction to date, a similar reporter gene approach successfully identified a transcriptional regulator of the sulfur deficiency response in Arabidopsis. This genetic screen used the same high-affinity sulfate transporter promoter element fused to the green fluorescent protein (*pSULTR1;2::GFP*) and identified four allelic mutants in an ethylene insensitive-like transcription factor called Sulfur Limitation 1 (SLIM1) that failed to induce the reporter construct under sulfur limiting conditions [14]. All of the allelic *slim1* mutants identified in this screen resulted in missense mutations altering single amino acid residues [14]. In *slim1-1* and *slim1-2*, high-affinity sulfate uptake was decreased by ~60%, and sulfur-dependent microarray analyses on *slim1-1* and *slim1-2* showed a decrease in the induction of many sulfur limitation-induced transcripts compared to controls suggesting that SLIM1 is a positive regulator of sulfate uptake and assimilation [14].

While the transcription factors that control arsenic-induced gene expression remain largely unknown, arsenic exposure is known to rapidly deplete cellular glutathione levels, increasing the demand for reduced sulfur compounds from the sulfur assimilation pathway[7,15,16]. A similar situation occurs under sulfur deficiency. As sulfate supply decreases, cellular levels of cysteine and glutathione become depleted. Thus, because of the similarities in glutathione depletion and subsequent upregulation of the high-affinity sulfate transporter *SULTR1;2* under arsenic stress [7] and sulfur limitation [14], we investigated the hypothesis that SLIM1 plays a role in arsenic-induced transcriptional responses. Interestingly, we found that *slim1-1* and *slim1-2* seedlings were highly sensitive to arsenic. Here, we show that under arsenic treatment, *slim1* mutants accumulate arsenic, experience high levels of oxidative stress, and fail to induce sulfate uptake and assimilation. Our results suggest that SLIM1 appears to play an important role in arsenic sensitivity due primarily to its role in regulating sulfur metabolism and the cellular redox state.

## Results

### *slim1* mutants are sensitive to arsenic in root growth assays

In a previous screen for regulators of cadmium and arsenic-induced gene expression using a *pSULTR1;2::LUC* reporter construct, we identified new alleles in well-characterized glutathione biosynthesis genes that play an essential role in cadmium and arsenic detoxification [7]. Because glutathione is a significant sink of reduced sulfur in plants, we hypothesized that the transcriptional regulator of sulfur deficiency, SLIM1, might also play a role in regulating cadmium and arsenic sensitivity in plants. To test this hypothesis, we performed root growth assays to evaluate the sensitivity of the *slim1-1* and *slim1-2* mutant alleles [14] to cadmium and arsenic (Figure 1A-1D).

**Figure 1.**
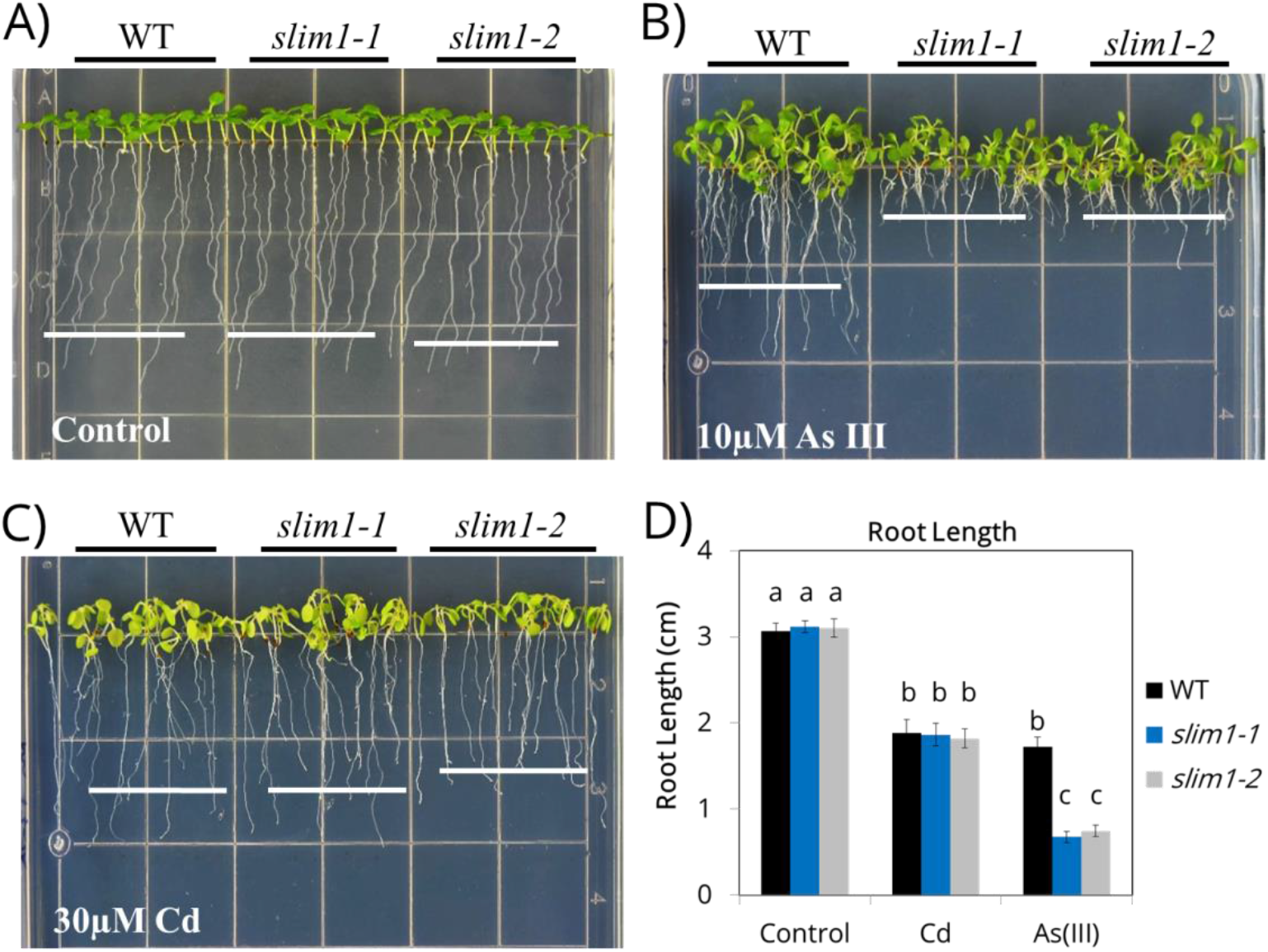
Root growth inhibition of *slim1* mutants grown on cadmium or arsenic-containing media. The *slim1-1* and *slim1-2* mutant alleles were compared to wild-type controls (WT) grown on control minimal media and media containing 30 μM Cd or 10 μM As(III) for 7 days (A - D). Root growth was quantified using ImageJ (one-way ANOVA, Tukey HSD).

The root lengths of wild-type (WT) (3.06 ± 0.09 cm, n=22), *slim1-1* (3.19 ± 0.07 cm, n=19), and *slim1-2* (3.10 ± 0.11 cm, n=21) were not different in the control nutrient media (see Methods) [17] without addition of cadmium or arsenic (Figure 1; p=0.99997 *slim1-1* & p=1.0 *slim1-2*, one-way ANOVA). When grown on plates containing 30 μM cadmium, WT root growth was inhibited growing only 1.88 ± 0.15 cm (n=10). This inhibition was similar to that observed for *slim1-1* with a final root length of 1.86 ± 0.13 cm (n=12) and *slim1-2* having a root length of 1.82 ± 0.11 cm (p=1.0 for *slim1-1* & p=0.99998 for *slim1-2*, n=13) (Figure 1A-1D, Table S1). However, when grown on minimal media plates containing 10 μM arsenite (As (III)), the root length of WT (1.72 ± 0.12 cm, n=14) was longer than both *slim1-1* (0.68 ± 0.06 cm, p=7×10-9, n=14) and *slim1-2* (0.75 ± 0.06 cm, p=1.2×10-6, n=10). These observations suggested that SLIM1 is involved in arsenic signaling. Thus, we further investigated possible mechanisms underlying *slim1* sensitivity to arsenic.

### Arsenic accumulation and antioxidant responses of *slim1* mutants

To determine if arsenic accumulates in the *slim1* mutants, we measured root and shoot arsenic levels using ICP-MS. In As(III) treated seedlings, we observed no significant increase in the accumulation of arsenic in the shoots of *slim1-1* (174.8 ± 1.97 mg/Kg DW, n=3) or *slim1-2* (189.7 ± 3.78 mg/Kg DW, n=3) compared to WT (173.9 ± 4.32 mg/Kg DW, n=3, p=1) (Figure 2A). However, in As(V) treated seedlings, both *slim1-1* (309.0 ± 47.5 mg/Kg DW, n=3, p=0.01) and *slim1-2* (253.9 ± 19.9 mg/Kg DW, n=3, p=0.6) accumulated more arsenic in shoots than WT (205.6 ± 11.6 mg/Kg DW, n=3), although the difference was only significant in *slim1-1* (Figure 2A). These results suggest an increased root-to-shoot translocation of As(V) in the *slim1* mutants.

**Figure 2.**
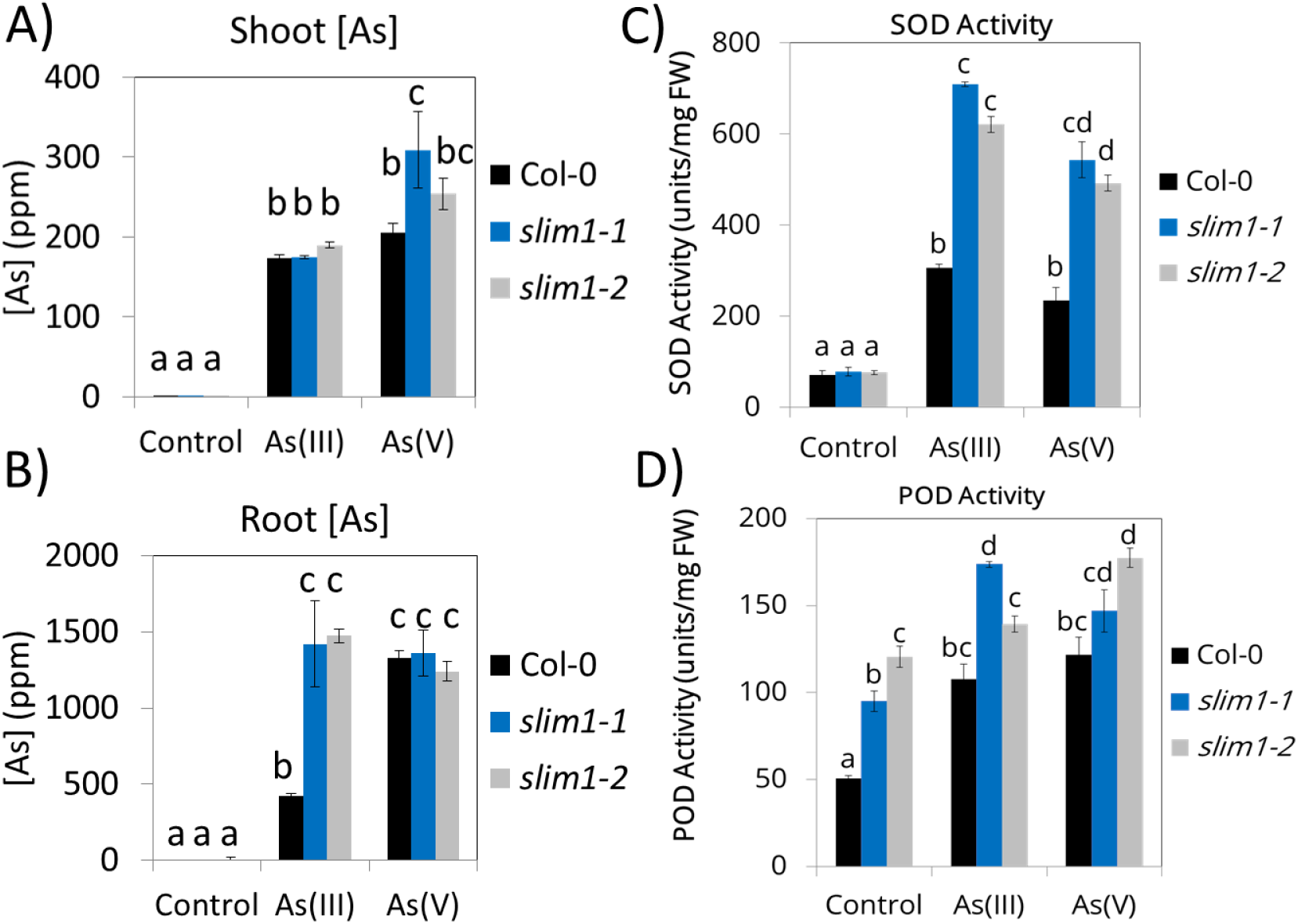
*slim1* mutants accumulate arsenic in roots and have high antioxidant activity when exposed to arsenic. *slim1* mutants grown on arsenic-containing media accumulate arsenic in the shoots when grown on As(V) (A) but accumulate arsenic in the roots when grown on As(III) (B). Growth on arsenic-containing media caused an increase in superoxide dismutase (C) enzyme and peroxidase dismutase enzyme (D) activities in both the *slim1-1* and *slim1-2* mutants compared to WT controls.

In the roots, we found arsenic accumulation in *slim1-1* (1420.0 ± 281.3 mg/Kg DW, n=3, p=6.77E-3) and *slim1-2* (1473.0 ± 187.9 mg/Kg DW, n=3, p=3.69E-3) compared to WT (420.1 ± 17.1 mg/Kg DW, n=3) in As(III) treated seedlings (Figure 2B, Table S3). In comparison, there was no difference in root arsenic accumulation in As(V) treated seedlings (Figure 2B, Table S3).

Because arsenic is known to cause oxidative stress and induce reactive oxygen species (ROS) production, we also tested the activity of the key antioxidant enzymes peroxidase (POD) and superoxide dismutase (SOD) in the *slim1-1* and *slim1-2* mutants. Basal superoxide dismutase activity in seedlings was similar between WT (70.9 ± 9.65 units/g FW, n=3), *slim1-1* (77.7 ± 9.44 units/g FW, p=1.0, n=3), and *slim1-2* (Figure 2C; 75.8 ± 4.20 units/g FW, p=1.0, n=3, p=1.0 *slim1-1* and p=1.0 *slim1-2*) grown under control conditions. When exposed to arsenite (As(III)), WT superoxide dismutase increased to 306.2 ± 7.78 units/g FW (n=3) while the superoxide dismutase activity in the *slim1* mutants increased dramatically to 709.5 ± 4.85 units/g FW (p=3.6×10-6, n=3) in *slim1-1* and 621.1 ± 17.7 units/g FW in *slim1-2* (p=2.0×10-8, n=3). Similarly, arsenate (As(V)) treatment increased the WT superoxide dismutase activity to 234.8 ± 27.2 units/g FW while the *slim1-1* superoxide dismutase activity increased to 543.2 ± 39.4 units/g FW (p=2.9×10-8, n=3) and the *slim1-2* superoxide dismutase activity increased to 492.1 ± 17.7 units/g FW (p=4.7×10-7, n=3) (Figure 2C, Table S4).

In seedlings, the peroxidase activity was higher under control conditions in *slim1-1* (94.8 ± 5.95 units/g FW, p=0.009, n=3) and *slim1-2* (120.6 ± 6.04 units/g FW, p=5.4×10-5, n=3) compared to WT (50.7 ± 1.51 units/g FW, n=3) (Figure 2D). As (III) exposure increased the peroxidase activity in WT to 107.5 ± 8.91 units/g FW (n=3) (Figure 2D), while the peroxidase activity in *slim1-1* seedlings increased to 173.6 ± 1.79 units/g FW (p=0.0001, n=3) (Figure 2D). Similar values were observed for As (III)-treated *slim1-2* seedlings (139.3 ± 4.49 units/g FW, p=0.10, n=3) (Figure 2D, Table S2). Peroxidase activities showed similar trends under As(V) treatment (Figure 2D, Table S5).

### Decreased shoot glutathione in arsenic-treated *slim1-1* and *slim1-2*

To determine if thiol production might also be altered by arsenic treatment in the *slim1* mutants, we measured root and shoot cysteine and glutathione levels using fluorescence HPLC of seedlings exposed to arsenite (As(III)) or arsenate (As(V)) for 48 hours (Figure 3A-3D). Shoot cysteine levels were lower in *slim1-1* (10.2 ± 0.5 pmol/mg FW, p=0.04, n=3) and *slim1-2* (12.2 ± 1.0 pmol/mg FW, 0.19, n=3) than WT (23.7 ± 2.7 pmol/mg FW, n=3) in control conditions (Figure 3A, Table S6). No clear decrease in the cysteine concentration was observed in response to As(III) or As(V) treatment (Figure 3A, Table S6).

**Figure 3.**
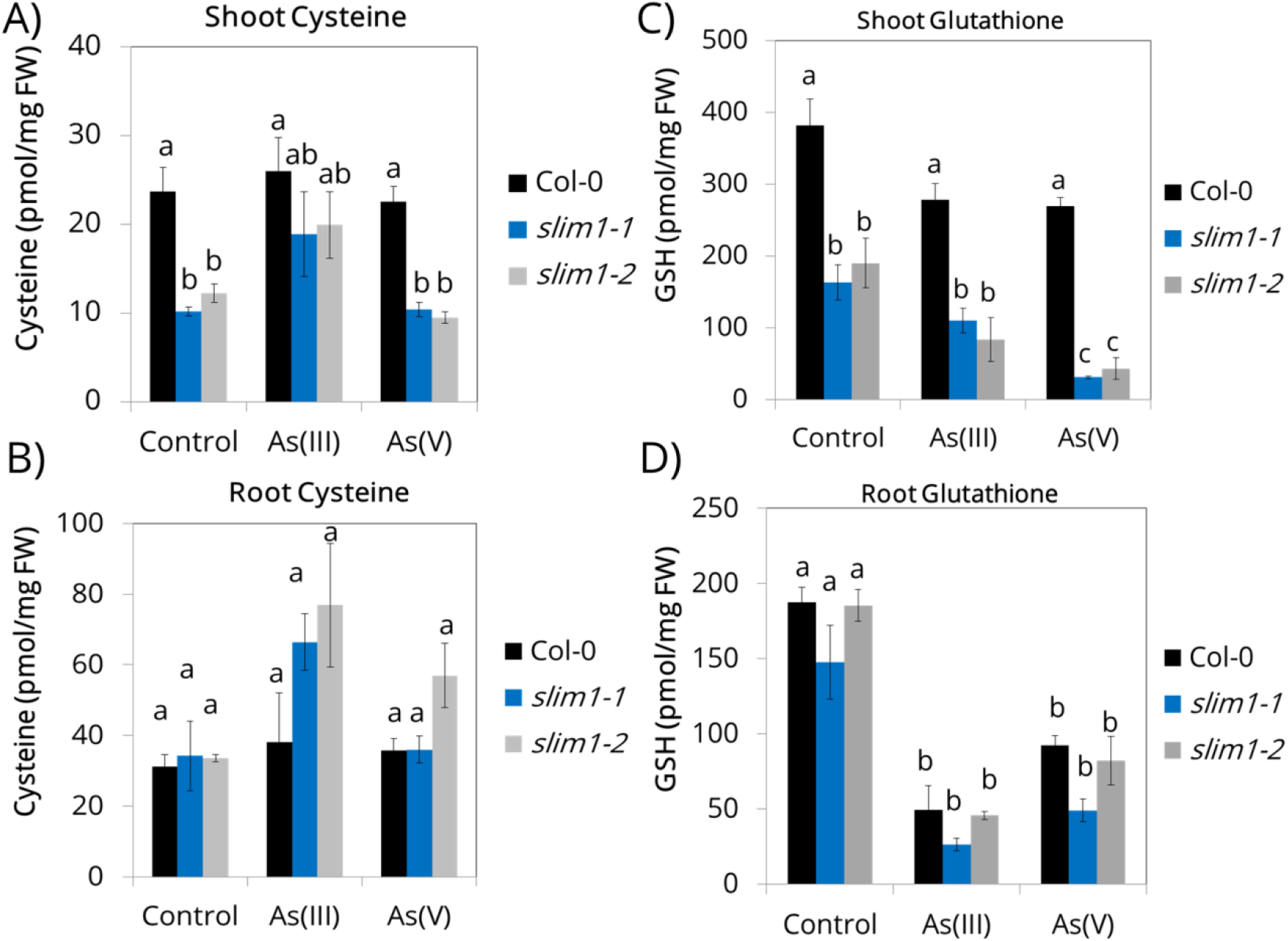
Thiol accumulation of *slim1* mutants grown on arsenic. Total shoot cysteine levels in *slim1-1* and *slim1-2* compared to WT (A). Total root cysteine levels in *slim1-1* and *slim1-2* compared to WT (B). Total shoot glutathione levels for *slim1-1* and *slim1-2* compared to WT (C). Total root glutathione levels for WT, *slim1-1*, and *slim1-2* (D).

Root cysteine levels were statistically similar for WT, *slim1-1*, and *slim1-2* in control conditions and were not significantly changed by As(III) or As(V) treatments (Figure 3B, Table S7; One-way ANOVA, Tukey HSD).

Under control conditions, shoot glutathione levels were lower in *slim1-1* (163.6 ± 24.2 pmol/mg FW, n=8) and *slim1-2* (190.2 ± 34.4 pmol/mg FW, n=8) than in WT (382.4 ± 36.2 pmol/mg FW, n=8) (Figure 3C; p=4.99×10-5 for *slim1-1* and p=3.9×10-4 for *slim1-2*). Shoot glutathione levels decreased in WT from 382.4 ± 36.2 pmol/mg FW (n=8) in control conditions to 278.1 ± 23.2 pmol/mg FW (n=3) in the As(III) treatment and 269.4 ± 12.2 pmol/mg FW (n=3) in the As(V) treatment (Figure 3C). Similarly, shoot glutathione decreased in the *slim1* mutants under As(III) and As(V) treatments with *slim1-1* having only 110.4 ± 17.5 pmol/mg FW of glutathione in As(III) and 31.4 ± 1.62 pmol/mg FW of glutathione in As(V). Furthermore, *slim1-2* had 83.8 ± 30.9 pmol/mg FW (n=3) shoot glutathione in As(III) treatment and 43.5 ± 14.7 pmol/mg FW (n=3) in As(V) treatment – an 80% decrease compared to control (Figure 3C, Table S8).

Root glutathione levels decreased under both As(III) and As(V) treatments for all genotypes. However, glutathione levels in roots showed no differences between genotypes within each treatment (Figure 3D, Table S9; One-way ANOVA, Tukey HSD). In summary, thiol measurements showed that while cysteine and glutathione levels were not dramatically decreased in the roots of the *slim1* mutant alleles compared to WT (Figure 3B and D), glutathione levels were decreased in shoots of *slim1-1* and *slim1-2* compared to WT plants (Figure 3C).

### Shoot sulfate and phosphate accumulation in *slim1* mutants

Arsenic is thought to be actively taken up by phosphate transporters as As(V); however, once inside plant cells, it is reduced to As(III) and can move within plants through aquaporins [18,19]. Mutants in SLIM1 were previously shown to be impaired in root-to-shoot translocation of sulfate [14], but the translocation of other anions, including phosphate, was not reported. Thus, based on the slight arsenic accumulation in shoots of As(V) treated plants noted by ICP-MS (Figure 2 A and B), we hypothesized that phosphate transport might also be impaired in the *slim1* mutants.

To determine if phosphate and sulfate translocation are impaired in the *slim1* mutants under arsenic treatment, we measured sulfate and phosphate accumulation in both roots and shoots of plants treated with As (V) for 48 hours. Interestingly, shoot phosphate accumulation was higher in *slim1-1* and *slim1-2* than WT in all treatments (Figure 4A, Table S10; p=5×10-6 for *slim1-1* and p=0.004 for *slim1-2*; One-way ANOVA, Tukey HSD).

**Figure 4.**
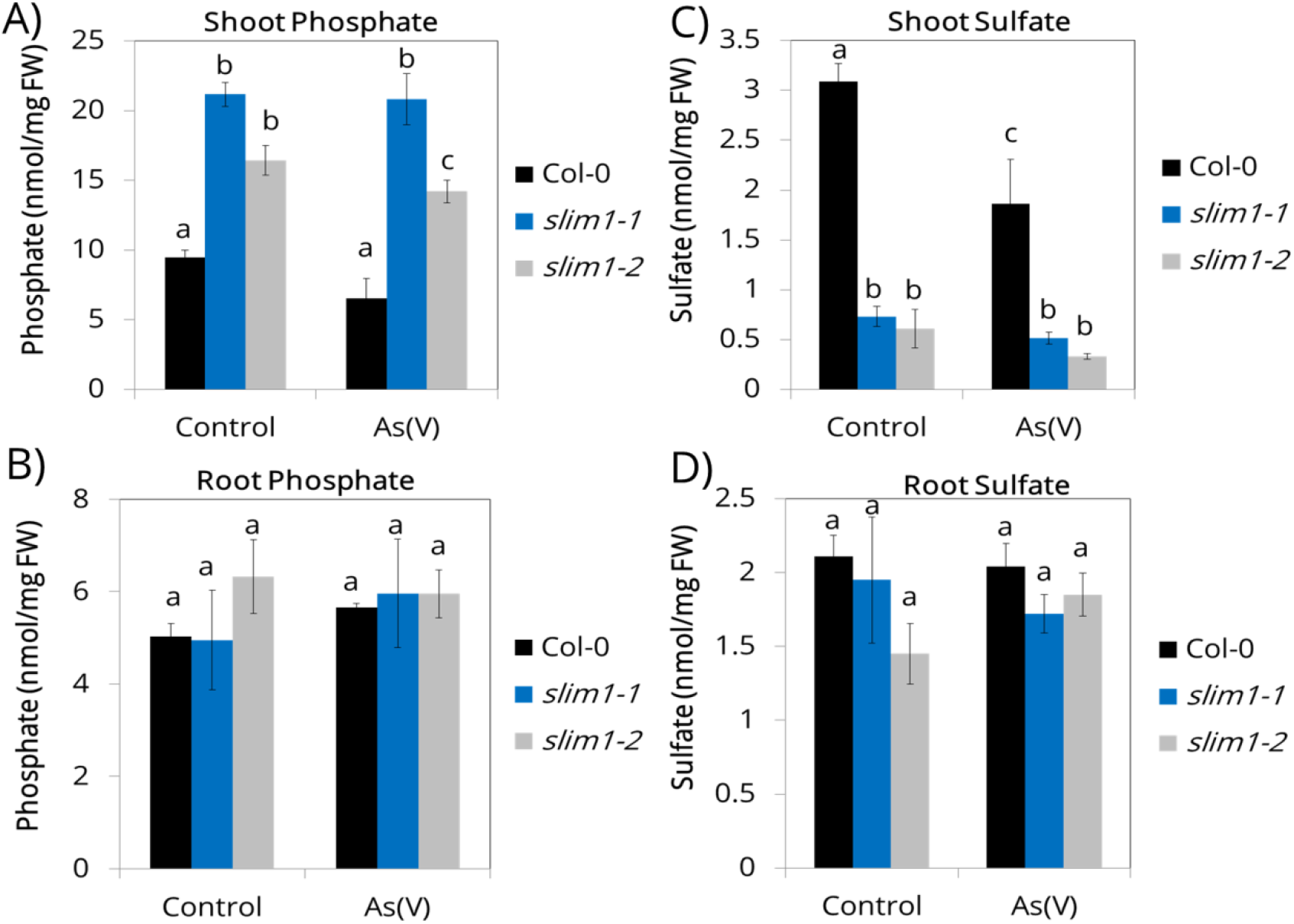
Anion accumulation in *slim1* mutants grown on arsenic. Total shoot phosphate levels in *slim1-1* and *slim1-2* compared to WT (A). Total root phosphate levels in *slim1-1* and *slim1-2* compared to WT (B). Total shoot sulfate levels for *slim1-1* and *slim1-2* compared to WT (C). Total root sulfate levels for WT, slim1-1, and *slim1-2* (D).

Root phosphate accumulation was similar for WT (5.03 ± 0.27 nmol/mg FW, n=5), *slim1-1* (4.95 ± 1.07 nmol/mg FW, n=3), and *slim1-2* (6.33 ± 0.80 nmol/mg FW, n=4) in control conditions and was not different under As(V) treatment (Figure 4B). Thus, the enhanced root-vs.-shoot phosphate accumulation observed in *slim1-1* and *slim1-2* suggests an indirect role for SLIM1 in regulating phosphate and arsenate transport (Figure 4A, 4B, Table S11).

Furthermore, sulfate accumulation in shoots was impaired in *slim1-1* (0.73 ± 0.10 nmol/mg FW, p=1.7×10-6, n=5) and *slim1-2* (0.61 ± 0.20 nmol/mg FW, p=7.3×10-7, n=5) relative to WT (3.09 ± 0.18 nmol/mg FW, n=5) in control conditions (Figure 4A), consistent with previous findings [14]. WT seedlings showed a decrease in shoot sulfate upon As(V) treatment decreasing to 1.86 ± 0.44 nmol/mg FW (n=5) (Figure 4C, Table S12, p=0.008, One-way ANOVA, Tukey HSD).

Root sulfate accumulation was similar between WT (2.11 ± 0.14 nmol/mg FW, n=5), *slim1-1* (1.95 ± 0.43 nmol/mg FW, n=3), and *slim1-2* (1.45 ± 0.21 nmol/mg FW, n=4) in control conditions. Furthermore, WT (2.04 ± 0.16 nmol/mg FW, n=4), *slim1-1* (1.72 ± 0.13 nmol/mg FW, n=5), and *slim1-2* (1.85 ± 0.15 nmol/mg FW, n=3) root sulfate were not different in the As(V) treatment (Figure 4D, Table S13).

### Microarray analyses of *slim1* mutants under As treatment

The current model for arsenic uptake and tolerance in plants suggests that arsenic is taken up from the soil in the form of arsenate (As(V)). Once it has entered the plant, it is rapidly reduced to arsenite (As(III)) by the arsenate reductase HAC1[20]. It has been proposed that As(III) can be removed from the root by an unidentified efflux transporter[21]. In rice, the aquaporin LSI1 is known to mediate As(III) efflux; however, additional efflux transporters remain elusive[21]. A recent RNA-seq experiment using a T-DNA mutant allele of SLIM1 (*eil3*) did not find misregulation of any aquaporin genes in the roots of the *slim1* mutant under control or sulfur deficiency conditions[22]. Thus, due to the observed arsenic accumulation in the roots of *slim1* mutants, we hypothesized that the elusive As(III) efflux transporter, or alternatively an As(III) uptake transporter, might be disrupted in an arsenic-dependent manner in the *slim1* mutant background.

To test this hypothesis and uncover genes disrupted in an arsenic-dependent manner in the *slim1-1* mutant, we performed microarray analyses on WT and *slim1-1* seedlings exposed to arsenite for 48 hours. Raw expression values were normalized via the R ‘affy’ package using the Robust Multi-Array Average (RMA) Expression Measure. Differential gene expression was evaluated using the R package ‘limma’, including a multiple test correction. We then performed a significance analysis to identify genes disrupted under arsenic treatment and compared these to previously published putative targets of SLIM1 obtained by DNA affinity purification sequencing (DAP-seq) [23].

From the microarray analyses, we identified 11 genes significantly differentially upregulated by arsenic (WT +As vs. *slim1-1* + As) (Supplemental Table S14). Ten of the 11 genes (AT3G49580, AT1G04770, AT1G12030, AT4G04610, AT4G21990, AT5G24660, AT5G26220, AT5G48850, AT4G20820, AT1G36370) were identified as putative targets of SLIM1 by DNA affinity purification sequencing (DAP-Seq) (Supplemental Table S14). Many of the 11 upregulated genes in *slim1-1* are associated with sulfur metabolism.

Genes that appear to be negatively regulated by SLIM1 include CGCT2;1 (AT5G26220), APR1 (AT4G04610) and APR3 (AT4G21990), which were upregulated in *slim1-1* compared to WT in the presence of arsenic (WT +As vs. *slim1-1* + As). APR1 and APR3 are involved in the reduction of sulfate into sulfide [24] and have been shown to be induced by toxic metal stress [7]. Similarly, the LOW SULFUR 1 (LSU1, AT3G49580) and LOW SULFUR 2 (LSU2, AT5G24660) genes were expressed at higher levels in *slim1-1* than WT under arsenic treatment (WT +As vs. *slim1-1* + As). Interestingly, Six of the 11 genes (GGCT2;1, APR3, LSU1, LSU2, SDI1, & SHM7) belong to a highly co-regulated cluster of genes that respond to O-acetylserine treatment[25].

Microarray analyses also identified 10 significantly down-regulated genes under arsenic treatment compared to WT (WT +As vs. *slim1-1* +As) (Supplemental Figure S1) (p<0.05, Fold Change >2). Only one gene - SULTR1; 2 (At1G78000) - was identified as a putative target of SLIM1 by DAP-Seq (Supplemental Table S14). Thus, our analyses confirm the reported function of SLIM1 as a transcriptional activator of *SULTR1;2* and show that this role is conserved under arsenic treatment and sulfur deficiency. The remaining ten genes are involved in hormone signaling (AT1G63030, AT5G13220 & AT5G52050), redox regulation (AT3G06590 & AT1G03020), iron homeostasis (AT3G25190 & AT5G01600), glucosinolate biosynthesis (AT5G23020), ubiquitination (AT1G24330), and an uncharacterized protein (AT2G17660). Based on their putative functions, these genes encode stress response-related genes. More experiments are needed to determine if SLIM1 is a direct transcriptional regulator of these genes under arsenic stress.

The present transcriptome data suggest that SLIM1 can function as both a transcriptional enhancer as well as a transcriptional repressor of specific genes in a condition-specific manner. Furthermore, the present study provides evidence that SLIM1 plays an essential role in the regulation of sulfur metabolism gene expression in response to arsenic.

## Discussion

Plant exposure to arsenic causes rapid changes in gene expression [7,26,27]. However, the transcription factors that function in arsenic-induced gene expression remain largely unknown. The few transcriptional regulators that have been identified, such as WRKY6, WRKY45, and OsARM1 (Arsenite-Responsive Myb1) [27–29], have been implicated in the regulation of arsenic transporters while regulators of arsenic detoxification remain unknown. To test the hypothesis that the SLIM1 transcription factor is involved in arsenic resistance and signaling, we evaluated the sensitivity of *slim1-1* and *slim1-2* to arsenic exposure. We found the *slim1* mutants were more sensitive to arsenic than control plants. Arsenic treatment caused high levels of oxidative stress in the *slim1* mutant alleles based on superoxide dismutase and peroxidase activities. Furthermore, thiol and sulfate measurements show that *slim1* mutants are impaired in both thiol and sulfate accumulation. Arsenic treatment did not further decrease sulfate levels in roots. In contrast, the concentration of the thiol GSH was greatly decreased in *slim1* mutant alleles. Furthermore, peroxidase and superoxide dismutase measurements show that arsenic treatments cause increased levels of oxidative stress in the *slim1* mutants.

We also observed a slight increase in arsenic accumulation in the shoots of *slim1* mutants treated with arsenic. This arsenic accumulation was accompanied by a significant increase in shoot phosphate translocation in the *slim1* mutants. Because of the chemical similarity between phosphate and arsenic oxyanions, future research could investigate the hypothesis that the misregulation of phosphate transporters may contribute to the observed increase in shoot arsenic in the *slim1* mutants. A recent study identified mutants in Ethylene Response Factor genes (ERF34 & ERF35) that are sensitive to both arsenite (As(III)) and arsenate (As(V)) [30]. Interestingly, similar to the *slim1* mutants, the double *erf34erf35* mutants were far less sensitive to cadmium than arsenic suggesting the arsenic sensitivity is not exclusively due to thiol accumulation. Furthermore, gene expression studies showed that several phosphate transporters were down-regulated in *erf34erf35* suggesting PHTs may play a role in both As(III) and As(V) sensitivity and/or transport [30].

Thiol measurements confirmed the role of SLIM1 in sulfate metabolism and thiol production [14,22,31], as *slim1* mutants contained lower cysteine and glutathione levels in shoots than WT. We hypothesized that the weaker cadmium sensitivity of *slim1* mutant alleles might be linked to thiol accumulation, but we observed no significant differences decrease in shoot GSH in the *slim1* mutants under Cd treatment (Supplemental Figure S2). However, SLIM1 upregulates the root-to-shoot transport of sulfate, which restricts sulfate assimilation mainly to the roots in *slim1* mutants. Root sulfate levels are maintained by the high-affinity sulfate transporter SULTR1;1, which is regulated in a SLIM1-independent manner [14]. Thus cysteine and glutathione biosynthesis can occur in the roots. As described previously, glutathione is essential for producing phytochelatins –arsenic chelating compounds necessary for detoxification and storage. The heavy metal cadmium also binds to phytochelatins. Interestingly, recent research has shown a less dramatic effect of cadmium exposure in *slim1* mutants than wild-type controls [31], which we have also observed (Figure 1C, D). Thus, the present study shows that the SLIM1 transcription factor plays a more central role in mediating arsenic resistance relative to cadmium resistance. A possible hypothesis that may contribute to this observation is that cadmium can be sequestered in vacuoles via two independent transport pathways: via phytochelatin transport [8,32] and via thiol-independent HMA3-mediated cadmium transport [33].

Sulfate measurements confirmed that SLIM1 is a major transcriptional regulator of sulfate uptake and translocation [14]. Our microarray analyses also identified 11 genes significantly differentially upregulated by arsenic (Supplemental Table S14), of which ten of the 11 genes were identified as putative targets of SLIM1 by DNA affinity purification sequencing (DAP-Seq). Interestingly, nine of these genes are involved in sulfur assimilation or redox signaling. One of these genes, GGCT2;1, is involved in glutathione recycling and has also been implicated in arsenic tolerance[34–36] Furthermore, six of these sulfur metabolism genes belong to a highly co-regulated cluster of genes that respond to O-acetylserine treatment [25]. While previous studies show these genes can regulate sulfur assimilation in a SLIM1 independent manner [25,37], results from DAP-Seq and microarray results from the current study suggest SLIM1 may act as a negative regulator of these genes during arsenic stress

Shoot sulfate accumulation was significantly lower in the *slim1* mutants under all conditions tested. Decreased shoot sulfate was accompanied by an increase in shoot phosphate in the *slim1* mutants. Similar anion compensation was noted in the Arabidopsis *phr1* mutant, which accumulates higher sulfate levels when grown under low phosphate conditions indicating crosstalk between phosphate and sulfate transport [38]. In fact, *PHR1* has been proposed to act both positively in the regulation of root-to-shoot sulfate translocation via the sulfate transporter *SULTR1;3*, and negatively to repress other sulfate transporters under phosphate deficiency [39]. We did not identify any significantly misregulated phosphate transporters (PHTs) in our microarray analyses. One possible explanation is that PHTs belong to a large gene family and demonstrate a high degree of genetic redundancy. Thus, a small decrease in the expression of several PHTs may result in measurable changes in phosphate accumulation without any individual transcript misregulation meeting the stringent criteria used in our microarray analyses. Xie et al. identified an artificial microRNA mutant targeting three high-affinity phosphate transporters showing a similar sensitivity to arsenite [30]. A recent study showing that sulfate deficiency increases phosphate accumulation in Arabidopsis further supports this hypothesis [40].

In summary, we show here that the SLIM1 transcription factor plays an important role in mediating arsenic resistance and in arsenic-induced gene expression. Our results suggest that the arsenic sensitivity of *slim1* mutants can be explained by decreased thiol production resulting in increased oxidative stress and in increased arsenic accumulation. Interestingly, we found that the *slim1* mutant alleles do not show a strong cadmium sensitivity, consistent with a recent study [31] indicating a difference in the rate-limiting functions of the thiol synthesis pathway in processing arsenic and cadmium that we discuss here. We also identify a number of genes regulated by SLIM1 in an arsenic-dependent manner with DAP-seq data set analyses indicating direct binding of SLIM1 to arsenic-dependent differentially-expressed genes. Taken together, our data support a model in which SLIM1 is both a positive and negative regulator of gene expression in response to arsenic.

## Experimental Procedures

### Arabidopsis accessions

The WT *Arabidopsis thaliana* ecotype used in this study is Columbia (Col-0). The *slim1-1* and *slim1-2* mutants were generated in the Col-0 genetic background and were kindly provided by Dr. Akiko Maruyama-Nakashita [14].

### Plant Growth Media & Conditions

Seeds were surface sterilized by briefly soaking in 70% ethanol before allowing them to dry in a sterile hood. For root growth experiments and enzymatic assay experiments, surface-sterilized seeds were plated on minimal media containing 1/10-strength Hoagland solution, 1% phytoagar (Duchefa, http://www.duchefa.com), pH 5.6. For the microarray experiments, seeds were plated on1/2-strength MS standard medium (M5519; Sigma-Aldrich, http://www.sigmaaldrich.com) buffered with 1 mM 2-(*N*-morpholine)-ethanesulphonic acid (MES), 1% phytoagar (Duchefa, http://www.duchefa.com) and the pH was adjusted to 5.6 with 1.0 m KOH. Seeds were then stratified with cold treatment at 4°C for 48 h, and grown under controlled conditions (150 μmol m^-2^ s^-1^, 70% humidity, 16-h light at 21°C/8-h dark at 18°C) for the specified time. For toxic metal(loid) treatments, the specified amounts of either cadmium or arsenic were added to the autoclaved base media in a sterile hood prior to pouring the plates. Concentrated stock solutions of cadmium and arsenic were filter-sterilized prior to use.

### Statistical Analyses

The root growth, thiol, peroxidase, superoxide, and anion data were all analyzed using one-way ANOVA followed by a Tukey posthoc test to determine significance. Significance groups are indicated in the figures, and key p-values are stated in the text.

### Length Measurements

For root growth experiments, surface-sterilized seeds of WT, *slim1-1*, and *slim1-2* were plated on minimal media (2.5 mM H_3_PO_4_, 5 mM KNO_3_, 2 mM MgSO_4_, 1 mM (CaNO_3_)_2_, 1 mM MES, 1% phytoagar pH 5.7) supplemented with 30 μM Cd or 10 μM As (III) [17]. Plates were placed in the dark two days at 4°C for vernalization and then transferred to a growth chamber. After 7 days of growth, seedlings were photographed, and root length was measured using ImageJ.

### Antioxidant Enzyme Assays

Seedling samples were weighed and pulverized in liquid nitrogen after treatment. The powder was dissolved in pre-cooled 50 mM phosphate buffer (pH 7.8) to extract the superoxide dismutase (SOD). The extract was then centrifuged at 12 000g for 10 min, resulting in a crude enzyme supernatant solution. In a separate 10 ml tube, 1.9 ml reaction buffer (50 mM phosphate buffer, pH 7.8, 9.9 mM L-methionine, 57 μM NBT solution, 1 M EDTA-Na_2_ solution, 0.0044% (w/v) riboflavin) and 0.1 ml enzyme solution were mixed and placed into 250 μmol m-2s-1 light for 20 min. Additionally, another separate 10 ml tube was procured, where the enzyme solution was replaced with water as a control. The reagent was added according to the above steps, where one tube was placed in the light together with the sample, and the other was placed in the dark where the reaction was allowed to complete. The control tube that was placed in the dark was blanked, and the absorbance of each tube was measured at 560 nm. Peroxidase (POD) was extracted in 50 mM phosphate buffer (pH 7.0). 30 μl of enzyme solution was mixed with reaction buffer containing 1.77 mL of 50 mM sodium phosphate buffer (pH 7.0), 0.1 mL of 4% guaiacol and 0.1 mL of 1% (v/v) H_2_O_2_. Increased absorbance was recorded at 470 nm for 1 min. All reported enzyme activities are means of 3-5 biologically independent samples, and error bars indicate the standard error of the mean (SEM).

### Arsenic Determination by ICP-MS

Plant material was harvested, dried at 70°C for at least 48 hours before being aliquoted and weighed. Approximately 10 mg of dried plant material was mixed with 1 ml of concentrated nitric acid and digested by heating at 100°C for approximately 30 minutes or until the solution became transparent and particle-free. These digests were diluted with deionized water and measured by ICP-MS for total arsenic concentrations at the University of Cologne Biocenter Mass Spectrometry Platform. All reported ion quantities are means of 3-5 biologically independent samples, and error bars indicate the standard error of the mean (SEM).

### Anion Extraction and Measurement by Ion Chromatography

To quantify the water-soluble anion concentrations (phosphate and sulfate) in plant tissues, 10-30 mg of fresh tissue was harvested and flash-frozen in liquid nitrogen. Frozen tissue was then pulverized using a bead mill (make & model), and anions were extracted by addition of 1000 μL of sterile Milli-Q-water and incubating for 60 minutes at 4°C while shaking at 1500 rpm. The extraction process was stopped by incubating at 95°C for 15 minutes. Cell debris was removed by centrifugation at 4°C for 15 minutes, and 100-200μL of supernatant was used for anion exchange chromatography. An automatic ion analyzer (DX 120, Dionex Corporation, Sunnyvale, CA, United States) equipped with an IonPac^TM^ column (AS9-SC, 4 × 250 mm; Dionex, Thermo Fisher Scientific GmbH; Waltham, MA, United States) was used to separate and quantify the anions. Anions were eluted with an elution buffer of 2.0 mM Na_2_CO_3_ and 0.75 mM NaHCO_3_. Ion concentrations were detected using a conductivity detector module (CDM, Dionex Corporation, CA, United States). All reported anion quantities are means of 3-5 biologically independent samples of tissue pooled from 4-6 individual seedlings (12-30 seedlings in total), and error bars indicate the standard error of the mean (SEM).

### Thiol Detection By Fluorescence HPLC

The thiol-containing compounds cysteine and GSH were analyzed using fluorescence detection HPLC as described by [41]. To analyze the levels of these thiol compounds, plants were grown on minimal growth media plates for 12 days then transferred to fresh media plates containing either 20 μM cadmium, 100 μM arsenate, or control minimal media. To minimize the oxidation of thiol compounds during the extraction, plant seedlings were flash-frozen in liquid nitrogen immediately after harvesting and then pulverized using a bead mill and extracted as described by [42]. Thiols were extracted from homogenized plant material with 1 mL 0.1 M HCl for 40 min at 25°C. After centrifugation for 5 min at 14,000 g and 4°C, thiols in the supernatant were reduced by mixing 60 μL of the supernatant with 100 μL 2-(cyclohexylamino)ethanesulfonic acid (0.25 M, pH 9.4) and 35 μL DTT (10 mM, freshly prepared). The mixture was incubated at 25°C for 40 min. Thiols were derivatized by adding 5 μL (25 mM) monobromobimane (SigmaAldrich, Cat#B4380). Derivatization was stopped by adding 110 μL methane sulfonic acid (100 mM) and clarified by centrifugation for 15 min at 14,000 g and 4°C.

Forty microliters of the derivatization mix were used for HPLC analysis using the Dionex Ultimate 3000 HPLC System. Derivatized thiols were separated in a Eurosphere 100-3 C18, 150×4 mm column (Knauer), and were detected by fluorescence detection with an excitation of 380 nm and emission detection at 480 nm. The peaks of thiol compounds were identified and quantified by comparison with cysteine and glutathione standards purchased from Sigma-Aldrich. All reported thiol quantities are statistical means of 4-5 biologically independent experiments (16-30 seedlings per experiment). Error bars indicate the standard error of the mean (SEM).

### Microarray Analyses

To evaluate transcriptional differences in the *slim1* mutants under cadmium and arsenic stress, we performed microarray analyses. To obtain tissue for the microarray analysis, plants were grown on ¼ MS plates for 12 days then transferred to fresh media plates containing either 100 μM cadmium or 20 μM arsenite. Whole seedlings were then harvested in 2mL Eppendorf tubes, flash-frozen in liquid nitrogen, and stored at −80°C until further processing. The tissue was subsequently pulverized using a bead mill by adding three 2.5mm glass beads to each tube and grinding for 15 seconds. RNA was extracted using the Qiagen RNEasy mini kit (Cat#74104) per the manufacturer’s instructions (www.qiagen.com). RNA quality was assessed by spectrophotometer and gel electrophoresis before submission to the University of California, San Diego Gene Expression Core facility for processing. Results were analyzed using R and the Bioconductor suite of microarray analytical packages as indicated in the text.

## Acknowledgments

We thank Dr. Akiko Maruyama-Nakashita for kindly providing *slim1* mutant seeds. This research was funded by the National Institute of Environmental Health Sciences of the National Institutes of Health under Award Number P42ES010337 (JIS). The microarray data are available through the NCBI GEO database (series record GSE138943).

## References

1 Clemens S (2006) Toxic metal accumulation, responses to exposure and mechanisms of tolerance in plants. Biochimie 88, 1707–1719.

2 Ogunseitan OA, Schoenung JM, Saphores J-DM & Shapiro AA (2009) The electronics revolution: from e-wonderland to e-wasteland. Science (80-) 326, 670–671.

3 Satarug S, Garrett SH, Sens MA & Sens DA (2009) Cadmium, environmental exposure, and health outcomes. Environ Health Perspect 118, 182–190.

4 Han FX, Banin A, Su Y, Monts DL, Plodinec JM, Kingery WL & Triplett GE (2002) Industrial age anthropogenic inputs of heavy metals into the pedosphere. Naturwissenschaften 89, 497–504.

5 Larison JR, Likens GE, Fitzpatrick JW & Crock JG (2000) Cadmium toxicity among wildlife in the Colorado Rocky Mountains. Nature 406, 181.

6 Mendoza-Cózatl DG, Xie Q, Akmakjian GZ, Jobe TO, Patel A, Stacey MG, Song L, Demoin DW, Jurisson SS, Stacey G & Schroeder JI (2014) OPT3 is a component of the iron-signaling network between leaves and roots and misregulation of OPT3 leads to an over-accumulation of cadmium in seeds. Mol Plant 7.

7 Jobe TO, Sung D, Akmakjian G, Pham A, Komives EA, Mendoza-Cózatl DG & Schroeder JI (2012) Feedback inhibition by thiols outranks glutathione depletion: a luciferase-based screen reveals glutathione-deficient γ-ECS and glutathione synthetase mutants impaired in cadmium-induced sulfate assimilation. Plant J 70, 783–795.

8 Mendoza-Cózatl DG, Zhai Z, Jobe TO, Akmakjian GZ, Song W-Y, Limbo O, Russell MR, Kozlovskyy VI, Martinoia E, Vatamaniuk OK, Russell P & Schroeder JI (2010) Tonoplast-localized Abc2 transporter mediates phytochelatin accumulation in vacuoles and confers cadmium tolerance. J Biol Chem 285.

9 Verbruggen N, Hermans C & Schat H (2009) Molecular mechanisms of metal hyperaccumulation in plants. New Phytol 181, 759–776.

10 Rea PA (2007) Plant ATP-binding cassette transporters. Annu Rev Plant Biol 58, 347–375.

11 Mendoza-Cózatl DG & Moreno-Sánchez R (2005) Cd2+ transport and storage in the chloroplast of Euglena gracilis. Biochim Biophys Acta (BBA)-Bioenergetics 1706, 88–97.

12 Jobe TO, Zenzen I, Rahimzadeh Karvansara P & Kopriva S (2019) Integration of sulfate assimilation with carbon and nitrogen metabolism in transition from C3 to C4 photosynthesis. J Exp Bot 70, 4211–4221.

13 Jobe, Timothy O., Kopriva S (2018) Sulfur Metabolism in Plants. Encycl Life Sci.

14 Maruyama-Nakashita A, Nakamura Y, Tohge T, Saito K & Takahashi H (2006) Arabidopsis SLIM1 is a central transcriptional regulator of plant sulfur response and metabolism. Plant Cell 18, 3235–3251.

15 Foyer CH & Noctor G (2011) Ascorbate and Glutathione: The Heart of the Redox Hub. Plant Physiol 155, 2 LP – 18.

16 Noctor G, Mhamdi A, Chaouch S, Han YI, Neukermans J, Marquez-Garcia B, Queval G & Foyer CH (2012) Glutathione in plants: an integrated overview. Plant Cell Environ 35, 454–484.

17 Lee DA, Chen A & Schroeder JI (2003) ars1, an Arabidopsis mutant exhibiting increased tolerance to arsenate and increased phosphate uptake. Plant J 35, 637–646.

18 Catarecha P, Segura MD, Franco-Zorrilla JM, García-Ponce B, Lanza M, Solano R, Paz-Ares J & Leyva A (2007) A mutant of the Arabidopsis phosphate transporter PHT1; 1 displays enhanced arsenic accumulation. Plant Cell 19, 1123–1133.

19 Bienert GP, Thorsen M, Schüssler MD, Nilsson HR, Wagner A, Tamás MJ & Jahn TP (2008) A subgroup of plant aquaporins facilitate the bi-directional diffusion of As (OH) 3 and Sb (OH) 3 across membranes. BMC Biol 6, 26.

20 Shi S, Wang T, Chen Z, Tang Z, Wu Z, Salt DE, Chao D-Y & Zhao F-J (2016) OsHAC1;1 and OsHAC1;2 Function as Arsenate Reductases and Regulate Arsenic Accumulation. Plant Physiol 172, 1708 – 1719.

21 Chen Y, Han Y-H, Cao Y, Zhu Y-G, Rathinasabapathi B & Ma LQ (2017) Arsenic Transport in Rice and Biological Solutions to Reduce Arsenic Risk from Rice. Front Plant Sci 8, 268.

22 Dietzen C, Koprivova A, Whitcomb SJ, Langen G, Jobe TO, Hoefgen R & Kopriva S (2020) The Transcription Factor EIL1 Participates in the Regulation of Sulfur-Deficiency Response. Plant Physiol 184, 2120 LP – 2136.

23 O’Malley RC, Huang SC, Song L, Lewsey MG, Bartlett A, Nery JR, Galli M, Gallavotti A & Ecker JR (2016) Cistrome and epicistrome features shape the regulatory DNA landscape. Cell 165, 1280–1292.

24 Lee B, Koprivova A & Kopriva S (2011) The key enzyme of sulfate assimilation, adenosine 5’-phosphosulfate reductase, is regulated by HY5 in Arabidopsis. Plant J 67, 1042–1054.

25 Hubberten H, Klie S, Caldana C, Degenkolbe T, Willmitzer L & Hoefgen R (2012) Additional role of O-acetylserine as a sulfur status-independent regulator during plant growth. Plant J 70, 666–677.

26 Fu S-F, Chen P-Y, Nguyen QTT, Huang L-Y, Zeng G-R, Huang T-L, Lin C-Y & Huang H-J (2014) Transcriptome profiling of genes and pathways associated with arsenic toxicity and tolerance in Arabidopsis. BMC Plant Biol 14, 94.

27 Castrillo G, Sánchez-Bermejo E, de Lorenzo L, Crevillén P, Fraile-Escanciano A, Tc M, Mouriz A, Catarecha P, Sobrino-Plata J, Olsson S, Leo del Puerto Y, Mateos I, Rojo E, Hernández LE, Jarillo JA, Piñeiro M, Paz-Ares J & Leyva A (2013) WRKY6 Transcription Factor Restricts Arsenate Uptake and Transposon Activation in <em>Arabidopsis</em> Plant Cell 25, 2944 LP – 2957.

28 Wang H, Xu Q, Kong Y-H, Chen Y, Duan J-Y, Wu W-H & Chen Y-F (2014) Arabidopsis WRKY45 Transcription Factor Activates <em>PHOSPHATE TRANSPORTER1;1</em> Expression in Response to Phosphate Starvation. Plant Physiol 164, 2020 LP – 2029.

29 Wang F-Z, Chen M-X, Yu L-J, Xie L-J, Yuan L-B, Qi H, Xiao M, Guo W, Chen Z, Yi K, Zhang J, Qiu R, Shu W, Xiao S & Chen Q-F (2017) OsARM1, an R2R3 MYB Transcription Factor, Is Involved in Regulation of the Response to Arsenic Stress in Rice. Front Plant Sci 8, 1868.

30 Xie Q, Yu Q, Jobe TO, Pham A, Ge C, Guo Q, Liu J, Liu H, Zhang H, Zhao Y, Xue S, Hauser F & Schroeder JI (2021) An amiRNA screen uncovers redundant CBF &amp; ERF34/35 transcription factors that differentially regulate arsenite and cadmium responses. bioRxiv.

31 Yamaguchi C, Khamsalath S, Takimoto Y, Suyama A, Mori Y, Ohkama-Ohtsu N & Maruyama-Nakashita A (2020) SLIM1 Transcription Factor Promotes Sulfate Uptake and Distribution to Shoot, Along with Phytochelatin Accumulation, Under Cadmium Stress in Arabidopsis thaliana. Plants 9, 163.

32 Song W-Y, Park J, Mendoza-Cózatl DG, Suter-Grotemeyer M, Shim D, Hörtensteiner S, Geisler M, Weder B, Rea PA & Rentsch D (2010) Arsenic tolerance in Arabidopsis is mediated by two ABCC-type phytochelatin transporters. Proc Natl Acad Sci 107, 21187–21192.

33 Takahashi R, Bashir K, Ishimaru Y, Nishizawa NK & Nakanishi H (2012) The role of heavy-metal ATPases, HMAs, in zinc and cadmium transport in rice. Plant Signal Behav 7, 1605–1607.

34 Joshi NC, Meyer AJ, Bangash SAK, Zheng Z-L & Leustek T (2019) Arabidopsis γ-glutamylcyclotransferase affects glutathione content and root system architecture during sulfur starvation. New Phytol 221, 1387–1397.

35 Paulose B, Chhikara S, Coomey J, Jung H, Vatamaniuk O & Dhankher OP (2013) A γ-Glutamyl Cyclotransferase Protects <em>Arabidopsis</em> Plants from Heavy Metal Toxicity by Recycling Glutamate to Maintain Glutathione Homeostasis. Plant Cell 25, 4580 LP – 4595.

36 Kumar S, Kaur A, Chattopadhyay B & Bachhawat AK (2015) Defining the cytosolic pathway of glutathione degradation in Arabidopsis thaliana: role of the ChaC/GCG family of γ-glutamyl cyclotransferases as glutathione-degrading enzymes and AtLAP1 as the Cys-Gly peptidase. Biochem J 468, 73–85.

37 Aarabi F, Naake T, Fernie AR & Hoefgen R (2020) Coordinating sulfur pools under sulfate deprivation. Trends Plant Sci.

38 Rouached H, Secco D, Arpat B & Poirier Y (2011) The transcription factor PHR1 plays a key role in the regulation of sulfate shoot-to-root flux upon phosphate starvation in Arabidopsis. BMC Plant Biol 11, 19.

39 Rouached H (2011) Multilevel coordination of phosphate and sulfate homeostasis in plants. Plant Signal Behav 6, 952–955.

40 Allahham A, Kanno S, Zhang L & Maruyama-Nakashita A (2020) Sulfur Deficiency Increases Phosphate Accumulation, Uptake, and Transport in Arabidopsis thaliana. Int J Mol Sci 21.

41 Fahey RC & Newton GL (1987) Determination of low-molecular-weight thiols using monobromobimane fluorescent labeling and high-performance liquid chromatography. In Methods in enzymology pp. 85–96. Elsevier.

42 Krueger S, Niehl A, Lopez Martin MC, Steinhauser D, Donath A, Hildebrandt T, Romero LC, Hoefgen R, Gotor C & Hesse H (2009) Analysis of cytosolic and plastidic serine acetyltransferase mutants and subcellular metabolite distributions suggests interplay of the cellular compartments for cysteine biosynthesis in Arabidopsis. Plant Cell Environ 32, 349–367.

