## Supplementary material for "The SLIM1 transcription factor is required for arsenic resistance in *Arabidopsis thaliana*": Supp_Data

**Supplementary Information:**

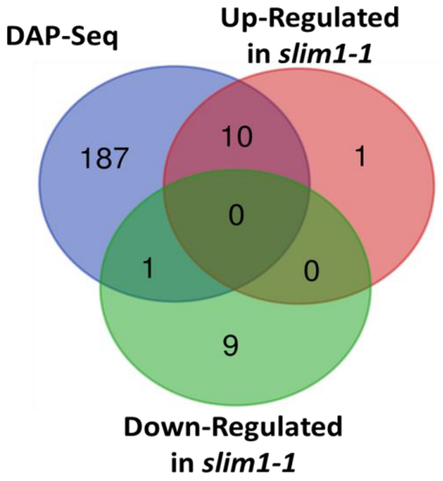


**Figure S1. Number of genes regulated by arsenic treatment compared with SLIM1 transcription factor DAP-Seq target genes.** Venn diagram showing the overlap of significantly upregulated genes in WT vs. *slim1-1* (pink) and down-regulated genes in WT vs. *slim1-1* (green) compared with previously published DAP-Seq targets of SLIM1 (blue). Gene lists can be found in Supplementary Table S14.

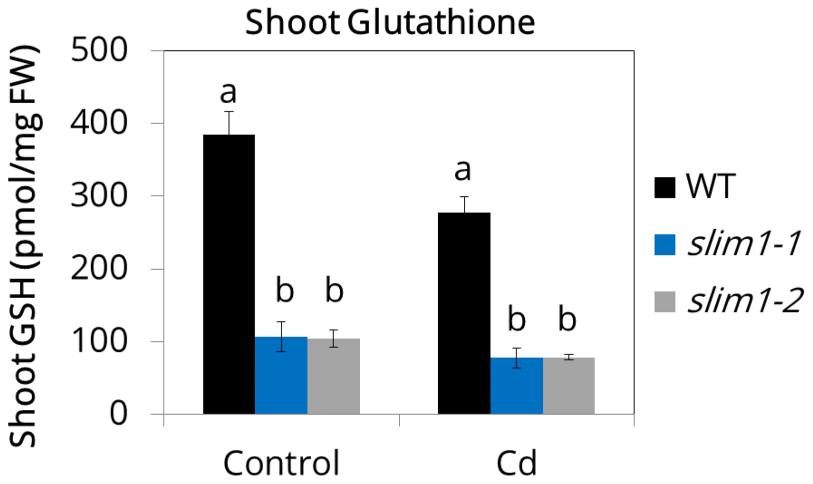


**Figure S2: Glutathione accumulation of *slim1* mutants grown on cadmium.** Total shoot glutathione levels for *slim1-1* and *slim1-2* compared to WT (ANOVA, Tukey HSD).

Table S1: Root Length On Cadmium And Arsenic

| Root Length (cm) | Control | | | Cd | | | As(III) | | |
|---|---|---|---|---|---|---|---|---|---|
| | WT | *slim1-1* | *slim1-2* | Col-0 | *slim1-1* | *slim1-2* | Col-0 | *slim1-1* | *slim1-2* |
| Average | 3.06 | 3.11 | 3.10 | 1.88 | 1.86 | 1.82 | 1.72 | 0.68 | 0.75 |
| N | 22 | 19 | 21 | 10 | 12 | 13 | 14 | 14 | 10 |
| Std Error | 0.09 | 0.06 | 0.11 | 0.15 | 0.13 | 0.11 | 0.11 | 0.06 | 0.06 |

Table S2: Shoot arsenic concentration (mg/Kg DW)

| Shoot Arsenic (mg/Kg DW) | Control | | | As(III) | | | As(V) | | |
|---|---|---|---|---|---|---|---|---|---|
| | WT | slim1-1 | slim1-2 | WT | slim1-1 | slim1-2 | WT | slim1-1 | slim1-2 |
| Avg | 0.24 | 0.20 | 0.18 | 173.9 | 174.8 | 189.7 | 205.6 | 309.0 | 253.9 |
| N | 3 | 3 | 3 | 3 | 3 | 3 | 3 | 3 | 3 |
| Std Error | 0.01 | 0.02 | 0.03 | 4.31 | 1.97 | 3.78 | 11.58 | 47.50 | 19.90 |

Table S3: Root arsenic concentration (mg/Kg DW)

| Root Arsenic (mg/Kg DW) | Control | | | As(III) | | | As(V) | | |
|---|---|---|---|---|---|---|---|---|---|
| | WT | slim1-1 | slim1-2 | WT | slim1-1 | slim1-2 | WT | slim1-1 | slim1-2 |
| Avg | 0.24 | 0.39 | 0.48 | 420.1 | 1420.0 | 1473.7 | 1331.2 | 1358.7 | 1241.1 |
| N | 3 | 3 | 3 | 3 | 3 | 3 | 3 | 3 | 3 |
| Std Error | 0.02 | 0.03 | 0.07 | 17.1 | 281.3 | 187.9 | 43.1 | 149.9 | 64.3 |

Table S4: SOD values under As(III) and As(V) treatment.

| SOD (units/mg FW) | Control | | | As(III) | | | As(V) | | |
|---|---|---|---|---|---|---|---|---|---|
| | WT | *slim1-1* | *slim1-2* | WT | *slim1-1* | *slim1-2* | WT | *slim1-1* | *slim1-2* |
| Average | 70.9 | 77.7 | 75.8 | 306.2 | 709.5 | 621.1 | 234.8 | 543.2 | 492.1 |
| N | 3 | 3 | 3 | 3 | 3 | 3 | 3 | 3 | 3 |
| Std Error | 9.65 | 9.45 | 4.20 | 7.78 | 4.85 | 17.72 | 27.22 | 39.44 | 17.69 |

Table S5: POD values under As(III) and As(V) treatment.

| POD (units/mg FW) | Control | | | As(III) | | | As(V) | | |
|---|---|---|---|---|---|---|---|---|---|
| | WT | *slim1-1* | *slim1-2* | WT | *slim1-1* | *slim1-2* | WT | *slim1-1* | *slim1-2* |
| Average | 50.73 | 94.82 | 120.61 | 107.51 | 173.61 | 139.30 | 121.54 | 146.94 | 177.44 |
| N | 3 | 3 | 3 | 3 | 3 | 3 | 3 | 3 | 3 |
| Std Error | 1.51 | 5.95 | 6.04 | 8.91 | 1.79 | 4.49 | 10.37 | 12.22 | 5.36 |

Table S6: Shoot Cysteine Levels Under Arsenic Treatment

| Shoot Cysteine | Control | | | As(III) | | | As(V) | | |
|---|---|---|---|---|---|---|---|---|---|
| (pmol/mg FW) | WT | *slim1-1* | *slim1-2* | Col-0 | *slim1-1* | *slim1-2* | Col-0 | *slim1-1* | *slim1-2* |
| Avg | 23.7 | 10.2 | 12.2 | 26.0 | 18.9 | 19.9 | 22.6 | 10.4 | 9.5 |
| N | 3 | 3 | 3 | 3 | 3 | 3 | 3 | 3 | 3 |
| Std Error | 2.7 | 0.5 | 1.0 | 3.8 | 4.8 | 3.7 | 1.7 | 0.8 | 0.6 |

Table S7: Root Cysteine Levels Under Arsenic Treatment

| Root Cysteine (pmol/mg FW) | Control | | | As(III) | | | As(V) | | |
|---|---|---|---|---|---|---|---|---|---|
| | WT | *slim1-1* | *slim1-2* | Col-0 | *slim1-1* | *slim1-2* | Col-0 | *slim1-1* | *slim1-2* |
| Avg | 31.2 | 34.2 | 33.6 | 38.1 | 66.4 | 76.8 | 35.7 | 35.9 | 56.9 |
| N | 3 | 3 | 3 | 3 | 3 | 3 | 3 | 3 | 3 |
| Std Error | 3.3 | 9.9 | 1.0 | 14.0 | 8.0 | 17.8 | 3.5 | 3.8 | 9.2 |

Table S8: Shoot Glutathione Levels Under Arsenic Treatment

| Shoot Glutathione (pmol/mg FW) | Control | | | As(III) | | | As(V) | | |
|---|---|---|---|---|---|---|---|---|---|
| | WT | *slim1-1* | *slim1-2* | Col-0 | *slim1-1* | *slim1-2* | Col-0 | *slim1-1* | *slim1-2* |
| Avg | 382.4 | 163.6 | 190.2 | 278.1 | 110.4 | 83.8 | 269.4 | 31.4 | 43.5 |
| N | 8 | 8 | 8 | 3 | 3 | 3 | 3 | 3 | 3 |
| Std Error | 36.2 | 24.2 | 34.4 | 23.2 | 17.5 | 30.9 | 12.2 | 1.6 | 14.7 |

Table S9: Root Glutathione Levels Under Arsenic Treatment

| Root Glutathione (pmol/mg FW) | Control | | | As(III) | | | As(V) | | |
|---|---|---|---|---|---|---|---|---|---|
| | WT | *slim1-1* | *slim1-2* | Col-0 | *slim1-1* | *slim1-2* | Col-0 | *slim1-1* | *slim1-2* |
| Avg | 187.4 | 147.7 | 185.5 | 49.5 | 26.4 | 45.6 | 92.1 | 49.1 | 82.1 |
| N | 3 | 3 | 3 | 3 | 3 | 3 | 3 | 3 | 3 |
| Std Error | 10.2 | 24.8 | 10.8 | 16.0 | 4.3 | 2.9 | 6.6 | 7.5 | 16.0 |

Table S10: Shoot phosphate values under As(V) treatment.

| Shoot Phosphate (pmol/mg FW) | Control | | | As(V) | | |
|---|---|---|---|---|---|---|
| | WT | *slim1-1* | *slim1-2* | WT | *slim1-1* | *slim1-2* |
| Avg | 9.5 | 21.2 | 16.4 | 6.5 | 20.8 | 14.2 |
| N | 5 | 5 | 5 | 5 | 5 | 4 |
| Std Error | 0.49 | 0.87 | 1.07 | 1.44 | 1.82 | 0.81 |

Table S11: Root phosphate values under As(V) treatment.

| Root Phosphate (pmol/mg FW) | Control | | | As(V) | | |
|---|---|---|---|---|---|---|
| | WT | *slim1-1* | *slim1-2* | WT | *slim1-1* | *slim1-2* |
| Avg | 5.03 | 4.95 | 6.33 | 5.66 | 5.96 | 5.95 |
| N | 5 | 3 | 4 | 4 | 5 | 3 |
| Std Error | 0.27 | 1.07 | 0.80 | 0.09 | 1.18 | 0.52 |

Table S12: Shoot sulfate values under As(V) treatment.

| Shoot Sulfate (pmol/mg FW) | Control | | | As(V) | | |
|---|---|---|---|---|---|---|
| | WT | *slim1-1* | *slim1-2* | WT | *slim1-1* | *slim1-2* |
| Avg | 3.09 | 0.73 | 0.61 | 1.86 | 0.52 | 0.33 |
| N | 5 | 5 | 5 | 5 | 5 | 4 |
| Std Error | 0.18 | 0.10 | 0.20 | 0.44 | 0.06 | 0.03 |

Table S13: Root sulfate values under As(V) treatment.

| Root Sulfate (pmol/mg FW) | Control | | | As(V) | | |
|---|---|---|---|---|---|---|
| | WT | *slim1-1* | *slim1-2* | WT | *slim1-1* | *slim1-2* |
| Avg | 2.11 | 1.95 | 1.45 | 2.04 | 1.72 | 1.85 |
| N | 5 | 3 | 4 | 4 | 5 | 3 |
| Std Error | 0.14 | 0.43 | 0.21 | 0.16 | 0.13 | 0.15 |
